# InSillyClo, a user-friendly web application to assist large-scale Golden Gate Cloning and MoClo workflows

**DOI:** 10.1101/2025.07.31.667881

**Authors:** Henri Galez, Bryan Brancotte, Juliette Bonche, Julien Fumey, Sara Napolitano, Gregory Batt

## Abstract

Systems and synthetic biology developments often require the construction of many variants of a genetic circuit of interest, resulting in large-scale cloning campaigns. Golden Gate and Modular Cloning (MoClo), two powerful technologies enabling the scale-up of cloning workflows, play a central role for efficient circuit construction. These workflows include a number of dry-lab tasks, which are time-consuming and error-prone at scale. Currently, no software tool is available to handle these tasks in a dedicated, time-saving, and user-friendly manner. We present InSillyClo, an open-source web application to assist large-scale Golden Gate cloning and MoClo workflows. It supports an easy specification of genetic designs at any scale, followed by the automated generation of comprehensive workflow-related data. Moreover, InSillyClo leverages Modular Cloning with a versatile typing system of parts to generate user-defined workflows. InSillyClo is open source, accessible with or without user registration, and can also be used locally.

## Introduction

Genetic engineering plays a pivotal role in synthetic and systems biology, either to empower living systems with new functions or to decipher their functioning. Despite decades of research works, engineering nature remains highly challenging. Design spaces have very high dimensionalities and knowledge of the functioning of cellular systems at the molecular level remains very partial and qualitative. In concrete terms, many variants of a target engineered system, or genetic circuit, have to be constructed to understand or optimize its functioning in its cellular context. This has motivated the development of a number of cloning technologies allowing efficient multipart, one-pot assemblies, such as Golden Gate^1^ and Gibson^2^ cloning. An extension of the Golden Gate assembly process is of special importance: Modular Cloning^3^ (MoClo). MoClo systems introduce standardized positions in the assembly (e.g., ‘part 2’, ‘part 3a’) and a biological significance to these positions (e.g., promoters are found as ‘part 2’). In computer science terms, MoClo systems are *typing* systems (e.g., promoters are of type ‘2’). This greatly simplifies the design of complex circuits. More than 50 MoClo kits are available on AddGene^4^, with some extending others with new parts and new part types, highlighting the interest of standardization.

Concretely, large cloning tasks are organized in cloning campaigns, in which one-pot assembly reactions are parallelized. These campaigns are typically composed of the following sequence of tasks: circuit design, assembly reaction, bacterial transformation, plasmid purification and verification (PCR, restriction profile and/or sequencing). Automation with liquid handling robots has significantly reduced the burden of the wet-lab part^5,6^. Yet, a number of tasks are conceptual in nature. The dry-lab part includes the specification of the genetic parts to assemble, the selection of the corresponding input plasmids, the computation of the final plasmid map, calculations for equimolar mixes based on part concentrations, and the computation of fragment length for PCR/restriction digestion for construct verification. These tasks are time-consuming and error-prone, especially for large cloning campaigns. Most software tools available for assisting Golden Gate cloning support only the manual design of plasmids, one at a time. Therefore, they are not well suited for large cloning campaigns. The few tools enabling multiple plasmids specification offer limited support for cloning campaigns or lack flexibility and synergy with MoClo.

Here, we present InSillyClo, a user-friendly web application to assist large-scale Golden Gate and modular cloning campaigns. InSillyClo offers a unique combination of features. It supports the specification of multiple assemblies at once with automated retrieval of input plasmid sequences from a laboratory data base, and the streamlined generation of valuable companion data, including plasmid maps, agarose gel simulation, dilution instructions and compatibility with automated cloning workflows. Moreover, InSillyClo leverages the notion of typing systems present in MoClo strategies to support campaign specification in a flexible and powerful manner. Lastly, InSillyClo is an open-source web application that can be used anonymously or as a logged user that offers confidentiality (no data stored for anonymous users) and user-friendliness (simulation history stored in user profiles). A command line version available as python package is also proposed for in-house, custom use.

## Results

### Support for large-scale Golden Gate Cloning campaigns

Typical cloning campaigns contain one or several combinatorial constructions together with a number of auxiliary modules. Here, we consider as a running example a combinatorial library to optimize the display of three cellulose degrading enzymes in yeast, together with the needed transcription factor, an optional anchor partner, and a stress reporter as auxiliary modules (Figure 1A).

**Figure 1:**
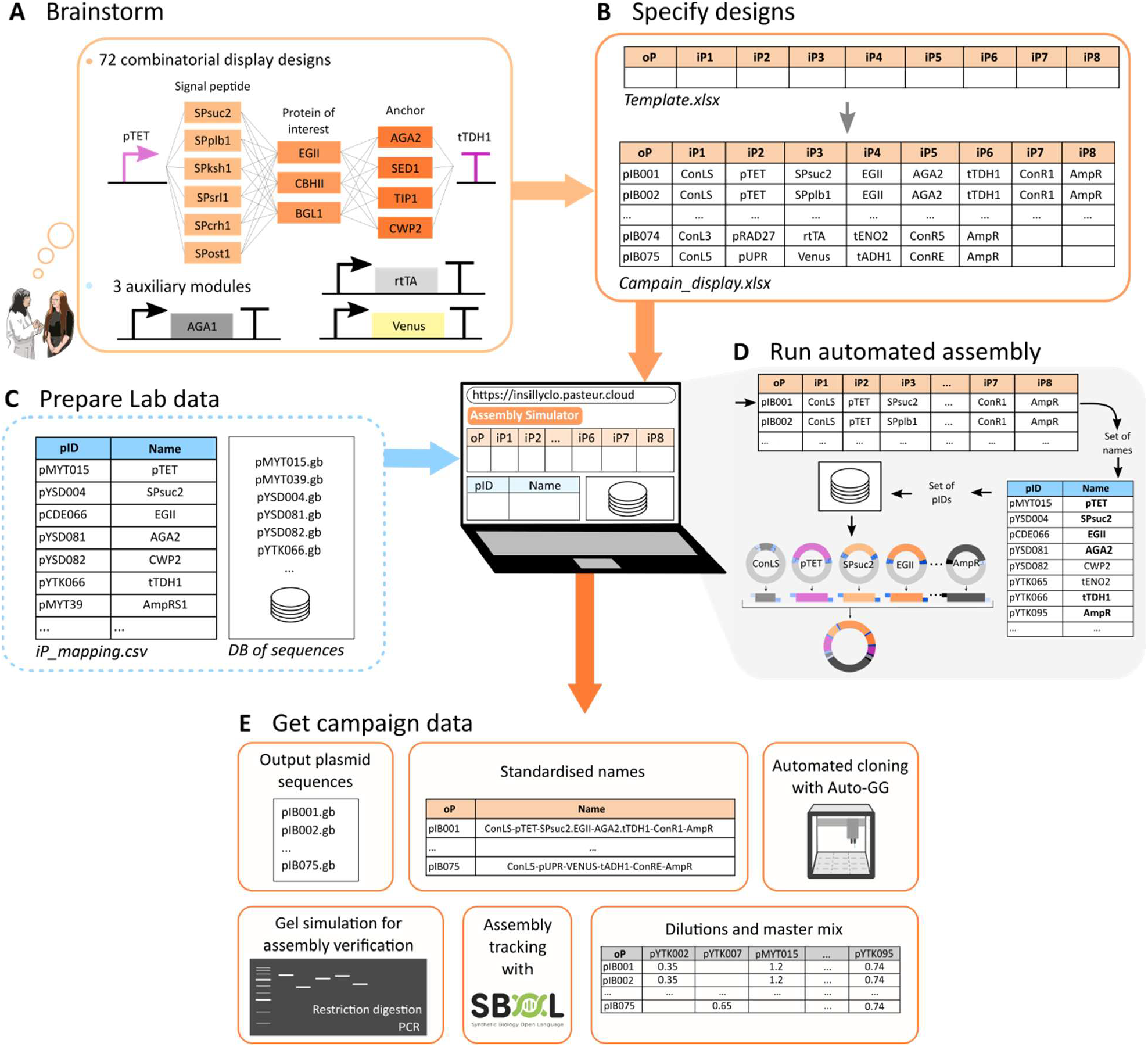
Golden Gate cloning campaigns with InSillyClo. A) Running example: designs for the inducible display of cellulose-degrading enzymes with various signal peptides and anchors. B) Design specification in a tabular format using a spreadsheet program using DNA part names. C) Upload of a database of input plasmids and of a mapping file for names. D) Algorithmic path from the scan of the cloning campaign file to the computation of output plasmid sequences. E) Workflow data generated by the Assembly Simulator module. Specifying primers and/or restriction enzyme is needed to generate simulated agarose gels. Filling in reaction parameters allows users to generate files for iP dilutions too.

These designs are specified by filling in a template file using a spreadsheet program (Figure 1B). Each line of the resulting *campaign* file represents a one-pot Golden Gate reaction. Specifications can be made using part names instead of plasmid identifiers (e.g. pTET instead of pMYT015; Text S1). This campaign file is loaded on the Assembly Simulator module of the web application, along with a database of Genbank files and one or several *iP_mapping* files (Figure 1C). These latter files are mapping part names with plasmid identifiers corresponding to Genbank file names. This InSillyClo specification process has been designed to combine simplicity, flexibility and efficiency. The Assembly Simulator algorithm performs the assemblies, one line at a time, by first gathering the part names, then retrieving the corresponding Genbank files through *iP_mapping*, and finally assembling the desired sequence (Figure 1D). The assembly is made by digesting input sequences with the chosen type IIS enzyme, retrieving the fragments inside the enzyme recognition sites, and joining them by matching overhangs.

The Assembly Simulator module will generate the sequences of all output plasmids, the *DB_produced* file, with an automated computation of output plasmid names for easy update of the *iP_mapping* database. It will also generate the instruction file for automated cloning with Opentrons liquid handling robots using the Auto-GG software tool^7^ (Figure 1E). Additional data can be generated depending on user needs. Automated computation of master mix and input plasmid dilutions is provided. Assistance for assembly verification is possible with the simulation of agarose gels after PCR and restriction digestion. An SBOL file containing the history of the assembly can be generated too.

InSillyClo was designed to assist experimentalists in their every-day lives by automating repetitive, time-consuming tasks. Interestingly, users will also get other benefits coming with automation, notably the automated computation of generic names and tables recapitulating mapping between names and plasmid identifiers. In that sense, InSillyClo can be seen as a minimal laboratory information management system, which could easily be integrated into more complete environments.

### Support for typed assemblies coming from MoClo systems

Modular Cloning approaches extend the Golden Gate technology by providing a standardization of the assemblies. In a given MoClo system, parts have specific positions in the assemblies. They are typed. One refers to them as parts of type 2, 3 or 4a for example. For each MoClo system, part types are specified by a particular combination of overhangs. The way parts assemble in constructs of different levels formally defines a grammar. This standardization considerably streamlines cloning processes. MoClo remains flexible thanks to the possibility of using spacers and subtyping. Modular Cloning has become instrumental in the genetic engineering of many organisms^8^. Figure 2A depicts the grammar of a popular MoClo system, the Yeast Tool Kit, which allows users to build transcriptional units and assemble them^9^.

**Figure 2:**
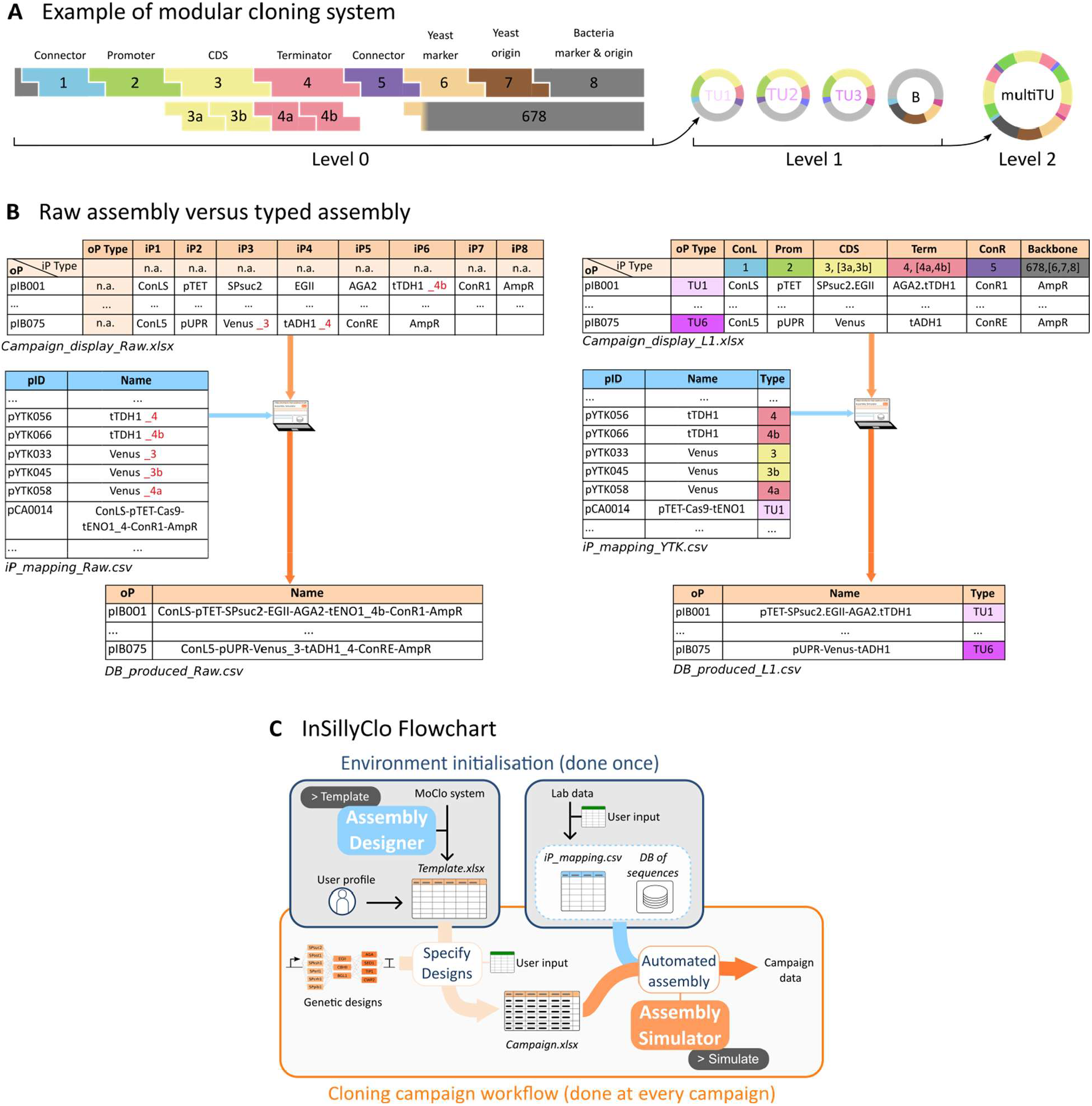
MoClo campaigns with typed assemblies using InSillyClo. A) Example of a MoClo grammar: the Yeast Tool Kit associates specific overhangs generated by Type IIs enzyme digestion to a biological meaning for the DNA parts. It forms a coherent assembly system to build (multi-)transcriptional units. B) Comparison between raw assembly (plain Golden Gate) and typed assembly (Moclo) for the campaign presented in Figure 1, underlying how the use of types clarifies the specifications. C) Overview of the full InSillyClo workflow, including the initialization phase.

InSillyClo takes advantage of the standardization offered by MoClo to provide a second mode for campaign specification that is more compact and clear. To illustrate the benefits at the design phase of MoClo over plain Golden Gate, we represent in Figure 2B the specification of the same cloning campaign with the plain Golden Gate format (raw assembly) or with the MoClo format (typed assembly). Several biological sequences are often present in part databases with different overhangs, that is, with different types, to be used at different positions. In our example, taken from the YTK and its extensions^10,11^, this is the case for the Venus reporter and the tTHD1 terminator. In the typed version, each part is represented by a *part-name / part-type* pair. In the raw version, the user needs to employ ad hoc name extensions to distinguish the otherwise identical parts, leading to error-prone situations. To embrace MoClo modularity, it is possible to define subparts by adding a *separator* character between two subparts as in the example with the dot “.” separating *AGA2* and *tTDH1*. Other advantages of the typed assembly specification are the design comfort with shorter standardized names and more homogeneous files.

Templates tailored to specific MoClo systems are generated with the Assembly Designer module of InSillyClo. The user can set the number of input parts, their generic names, the possibility to have subparts, the associated type IIs enzyme, and the naming conventions. Templates are already available on the Web App for widely-used MoClo kits.

New users need to perform the initialization step (Figure 2C). It involves downloading or generating the template (Text S2), preparing the repository of plasmid sequences, and providing the file mapping part names and types with plasmid unique identifiers (*iP_mapping* file). Templates for several widely-used MoClo systems (YTK^9^, Plant Part Kit^12^, Cidar^13^, and EcoFlex^14^) are available in the InSillyClo Gitlab repository (Text S3). Then, for each campaign, the user will fill in the template thus obtaining the *campaign* file. Running the Assembly Simulator module will generate the campaign data. InSillyClo is also available as a command line tool in which the *template* and *simulate* commands mirror the Assembly Designer and Assembly Simulator Web App modules (Text S4). The tool consists in a python package available in Pypi.

## Discussion

This work presents InSillyClo, an open-source web application supporting Golden Gate Cloning and Moclo workflows. It supports simple and flexible specification of genetic designs, even for large-scale campaigns, with automated retrieval of input sequences and assembly of output sequences. In addition, the application generates data helping users in wet lab tasks and in reporting. InSillyClo embraces the typing system present in MoClo to improve user experience. The tool is free of charge and runs on servers hosted by the Institut Pasteur, a non-profit organization. It is possible to use the web application anonymously (without registration). Alternatively, a command line tool containing most features can be used locally. Tutorials are available on the web application for a smooth introduction, and templates for many MoClo kits are already provided.

Golden Gate Cloning is supported by many software tools. Several tools guide users for manual plasmid construction with an intuitive interface (Ape^15^, OpenCloning^16^, SnapGene^17^, Benchling^18^, Geneious^19^, Teselagen^20^). With such tools, plasmids are built one at a time. This approach is therefore not suitable for large cloning campaigns. Some tools allow users to specify multiple plasmids to be constructed at the same time. This is notably the case for Benchling^18^, Geneious^19^, Teselagen^20^, Diva^21^ and Cuba^22^. Cuba^22^ is the only one in which the retrieval of input sequences is automated, as in InSillyClo, which is important for large campaigns. The specification phase is done in a tabular format for both Cuba^22^ and InSillyClo. However, InSillyClo users can refer to input plasmids directly with part names, whereas Cuba users need to fetch the corresponding Genbank file name or the Genbank internal ID field. While many of the tools presented above provide calculations for equimolar reaction mixes based on part concentrations and/or support assembly verification via simulation of agarose gels after PCR and restriction digestion, InSillyClo streamlines data generation thanks to its campaign-oriented approach. This approach differs from other pipelines dedicated to high-throughput Golden Gate cloning combining several software tools and modules^23^. In addition, InSillyClo uniquely supports MoClo by leveraging its typing system and letting users tailor the workflow to their MoClo system. Several improvements are envisioned. They notably include the support of Golden Gate assemblies with two restriction enzymes, the integration of genetic part homing by PCR, the connection with popular databases, and the compliance of the generated SBOL files with a recent standard^24^.

Large biofoundries are equipped with comprehensive software ecosystems as demonstrated by the applications developed by the Edinburgh Genome Foundry^23,25–28^. Yet, automation of Golden Gate and MoClo workflows is amenable for small companies or academic laboratories, via the standardization of manual protocols or the use of liquid handling robots, increasingly accessible^5,7^. We therefore believe that by providing support for the design, verification, reporting, and wet-lab automation, InSillyClo will usefully serve the bioengineering and synthetic biology communities.

## Methods

InSillyClo is distributed under the LGPL-3 license. It is organized as a python command line tool wrapped in a web application. The command line tool performs the computations and produces the data. It can also be used independently as a python package installable by any package manager since it is available on Pypi (Text S4). InSillyClo leverages biopython for handling of sequence files, pandas for template parsing, and pycairo for image generation. The web application is built on Django, Bootstrap, Docker and is hosted on a kubernetes cluster at Institut Pasteur. The web application guides users in the generation of their assembly template, then simulates Golden Gate assemblies of plasmid designs specified in the template.

## Associated content

### Data availability statement

InSillyClo can be accessed at https://insillyclo.pasteur.cloud. The source code is available at https://gitlab.pasteur.fr/insillyclo. Documentation for the Web App is available in the Tutorial section of the application. Documentation for the command line tool is available at https://insillyclo.pages.pasteur.fr/insillyclo-cli/.

### Supporting information

Supplementary information is available as a companion file.

## Author information

### Author contributions

H.G. proposed the project and developed prototype code.

H.G. and G.B. co-supervised the project, with help from S.N..

J.B. designed the software font-end.

B.B. developed the back-end and implemented the front-end of the software, with help from J.F..

H.G. and G.B. wrote the paper.

### Funding

This work was supported by ANR grants Anoruti (ANR-20-PAMR-0001), Seq2Diag (ANR-20-PAMR-0010), SmartSec (ANR-21-CE44-0033) and TrojanYeast (ANR-24-CE18-2885).

## Acknowledgments

We want to thank the Biofoundry team from Lesaffre, notably François Bertaux. Our collaboration led to the idea of developing InSillyClo software. We also thank Esteban Lebrun and Alicia Da Silva, who tested early versions of the software and provided useful feedbacks. Lastly, we thank Rachel Torchet who helped for the front-end design.

## Abbreviations

MoClo: Modular Cloning
YTK: Yeast Tool Kit
TU: Transcriptional Unit
iP: input Plasmid
oP: output Plasmid
SBOL: Synthetic Biology Open Language

## Notes

https://insillyclo.pasteur.cloud

https://gitlab.pasteur.fr/insillyclo

### Competing Interest Statement

The authors have declared no competing interest.

## References

(1) Engler, C.; Gruetzner, R.; Kandzia, R.; Marillonnet, S. Golden Gate Shuffling: A One-Pot DNA Shuffling Method Based on Type IIs Restriction Enzymes. PLOS ONE 2009, 4 (5), e5553. 10.1371/journal.pone.0005553.

(2) Gibson, D. G.; Young, L.; Chuang, R.-Y.; Venter, J. C.; Hutchison, C. A.; Smith, H. O. Enzymatic Assembly of DNA Molecules up to Several Hundred Kilobases. Nat. Methods 2009, 6 (5), 343–345. 10.1038/nmeth.1318.

(3) Weber, E.; Engler, C.; Gruetzner, R.; Werner, S.; Marillonnet, S. A Modular Cloning System for Standardized Assembly of Multigene Constructs. PLOS ONE 2011, 6 (2), e16765. 10.1371/journal.pone.0016765.

(4) Addgene: Modular Cloning Guide. https://www.addgene.org/cloning/moclo/(accessed 2025-07-24).

(5) Bryant, J. A.; Wright, R. C. Biofoundry-Assisted Golden Gate Cloning with AssemblyTron. In Golden Gate Cloning: Methods and Protocols; Schindler, D., Ed.; Springer US: New York, NY, 2025; pp 133– 147. 10.1007/978-1-0716-4220-7_8.

(6) Kang, D. H.; Ko, S. C.; Heo, Y. B.; Lee, H. J.; Woo, H. M. RoboMoClo: A Robotics-Assisted Modular Cloning Framework for Multiple Gene Assembly in Biofoundry. ACS Synth. Biol. 2022, 11 (3), 1336– 1348. 10.1021/acssynbio.1c00628.

(7) Meng, F.; Malci, K. Auto-GG, 2024. https://github.com/FankangMeng/Auto-GG (accessed 2025-07-11).

(8) Bird, J. E.; Marles-Wright, J.; Giachino, A. A User’s Guide to Golden Gate Cloning Methods and Standards. ACS Synth. Biol. 2022, 11 (11), 3551–3563. 10.1021/acssynbio.2c00355.

(9) Lee, M. E.; DeLoache, W. C.; Cervantes, B.; Dueber, J. E. A Highly Characterized Yeast Toolkit for Modular, Multipart Assembly. ACS Synth. Biol. 2015, 4 (9), 975–986. 10.1021/sb500366v.

(10) Shaw, W. M.; Khalil, A. S.; Ellis, T. A Multiplex MoClo Toolkit for Extensive and Flexible Engineering of Saccharomyces Cerevisiae. ACS Synth. Biol. 2023, 12 (11), 3393–3405. 10.1021/acssynbio.3c00423.

(11) O’Riordan, N. M.; Jurić, V.; O’Neill, S. K.; Roche, A. P.; Young, P. W. A Yeast Modular Cloning (MoClo) Toolkit Expansion for Optimization of Heterologous Protein Secretion and Surface Display in Saccharomyces Cerevisiae. ACS Synth. Biol. 2024. 10.1021/acssynbio.3c00743.

(12) Engler, C.; Youles, M.; Gruetzner, R.; Ehnert, T.-M.; Werner, S.; Jones, J. D. G.; Patron, N. J.; Marillonnet, S. A Golden Gate Modular Cloning Toolbox for Plants. ACS Synth. Biol. 2014, 3 (11), 839– 843. 10.1021/sb4001504.

(13) Iverson, S. V.; Haddock, T. L.; Beal, J.; Densmore, D. M. CIDAR MoClo: Improved MoClo Assembly Standard and New E. Coli Part Library Enable Rapid Combinatorial Design for Synthetic and Traditional Biology. ACS Synth. Biol. 2016, 5 (1), 99–103. 10.1021/acssynbio.5b00124.

(14) Moore, S. J.; Lai, H.-E.; Kelwick, R. J. R.; Chee, S. M.; Bell, D. J.; Polizzi, K. M.; Freemont, P. S. EcoFlex: A Multifunctional MoClo Kit for E. Coli Synthetic Biology. ACS Synth. Biol. 2016, 5 (10), 1059–1069. 10.1021/acssynbio.6b00031.

(15) Davis, M. W.; Jorgensen, E. M. Using ApE for In Silico Golden Gate Cloning. In Golden Gate Cloning: Methods and Protocols; Schindler, D., Ed.; Springer US: New York, NY, 2025; pp 79–87. 10.1007/978-1-0716-4220-7_5.

(16) OpenCloning. https://opencloning.org/(accessed 2025-07-11).

(17) SnapGene Software. https://www.snapgene.com/ (accessed 2025-07-09).

(18) Benchling Biology Software. https://www.benchling.com (accessed 2025-07-09).

(19) Geneious Prime. Geneious. https://www.geneious.com/series/prime (accessed 2025-07-11).

(20) Teselagen. https://teselagen.com/(accessed 2025-07-11).

(21) DIVA bioCAD. https://public-diva.agilebiofoundry.org/login (accessed 2025-07-11).

(22) Zulkower, V. Computer-Aided Design and Pre-Validation of Large Batches of DNA Assemblies. In Synthetic Gene Circuits: Methods and Protocols; Menolascina, F., Ed.; Springer US: New York, NY, 2021; pp 157–166. 10.1007/978-1-0716-1032-9_6.

(23) Vegh, P.; Chapman, E.; Gilmour, C.; Fragkoudis, R. Modular DNA Construct Design for High-Throughput Golden Gate Assembly. In Golden Gate Cloning: Methods and Protocols; Schindler, D., Ed.; Springer US: New York, NY, 2025; pp 61–77. 10.1007/978-1-0716-4220-7_4.

(24) Vidal, G.; Vitalis, C.; Guillén, J. Standardized Golden Gate Assembly Metadata Representation Using SBOL. In Golden Gate Cloning: Methods and Protocols; Schindler, D., Ed.; Springer US: New York, NY, 2025; pp 89–104. 10.1007/978-1-0716-4220-7_6.

(25) Hillson, N.; Caddick, M.; Cai, Y.; Carrasco, J. A.; Chang, M. W.; Curach, N. C.; Bell, D. J.; Le Feuvre, R.; Friedman, D. C.; Fu, X.; Gold, N. D.; Herrgård, M. J.; Holowko, M. B.; Johnson, J. R.; Johnson, R. A.; Keasling, J. D.; Kitney, R. I.; Kondo, A.; Liu, C.; Martin, V. J. J.; Menolascina, F.; Ogino, C.; Patron, N. J.; Pavan, M.; Poh, C. L.; Pretorius, I. S.; Rosser, S. J.; Scrutton, N. S.; Storch, M.; Tekotte, H.; Travnik, E.; Vickers, C. E.; Yew, W. S.; Yuan, Y.; Zhao, H.; Freemont, P. S. Building a Global Alliance of Biofoundries. Nat. Commun. 2019, 10 (1), 2040. 10.1038/s41467-019-10079-2.

(26) Zulkower, V.; Rosser, S. DNA Chisel, a Versatile Sequence Optimizer. Bioinformatics 2020, 36 (16), 4508–4509. 10.1093/bioinformatics/btaa558.

(27) Doçi, G.; Fuchs, L.; Kharbanda, Y.; Schickling, P.; Zulkower, V.; Hillson, N.; Oberortner, E.; Swainston, N.; Kabisch, J. DNA Scanner: A Web Application for Comparing DNA Synthesis Feasibility, Price and Turnaround Time across Vendors. Synth. Biol. 2020, 5 (1), ysaa011. 10.1093/synbio/ysaa011.

(28) Zulkower, V.; Rosser, S. DNA Features Viewer: A Sequence Annotation Formatting and Plotting Library for Python. Bioinformatics 2020, 36 (15), 4350–4352. 10.1093/bioinformatics/btaa213.

(29) Melero-Cobo, X.; Gallemí, M.; Carnicer, M.; Monte, E.; Planas, A.; Leivar, P. MoCloro: An Extension of the Chlamydomonas Reinhardtii Modular Cloning Toolkit for Microalgal Chloroplast Engineering. Physiol. Plant. 2025, 177 (1), e70088. 10.1111/ppl.70088.

(30) Harmer, Z. P.; McClean, M. N. The Yeast Optogenetic Toolkit (yOTK) for Spatiotemporal Control of Gene Expression in Budding Yeast. In Optogenetics: Methods and Protocols; Baumschlager, A., Ed.; Springer US: New York, NY, 2025; pp 19–36. 10.1007/978-1-0716-4047-0_2.

(31) Koh, H. G.; Tohidifar, P.; Oh, H.; Ye, Q.; Jung, S.-C.; Rao, C. V.; Jin, Y.-S. RT-EZ: A Golden Gate Assembly Toolkit for Streamlined Genetic Engineering of Rhodotorula Toruloides. ACS Synth. Biol. 2025, 14 (5), 1572–1580. 10.1021/acssynbio.4c00848.

(32) Soltysiak, M. P. M.; Ory, A. L. H.; Lee, A. D.; Christophersen, C. E.; Jalihal, A. P.; Springer, M. XanthoMoClo─A Robust Modular Cloning Genetic Toolkit for the Genera Xanthobacter and Roseixanthobacter. ACS Synth. Biol. 2025, 14 (4), 1173–1190. 10.1021/acssynbio.4c00806.

