## Supplementary Text for "InSillyClo, a user-friendly web application to assist large-scale Golden Gate Cloning and MoClo workflows"

### Summary

Text S1: Quick cloning with InSillyClo

Text S2: Standard typed cloning workflow

Text S3: InSillyClo with other MoClo systems

Text S4: Local implementation with the command line tool

File S1: Campaign_display_Raw.xlsx

File S2: iP_mapping_Raw.csv

File S3: Campaign_display_L1.xlsx

File S4: iP_mapping_YTK.csv

### Text S1: Quick cloning with InSillyClo

For users primarily interested in the automation of Golden Gate reactions, a minimal version of the workflow is possible:

1. Download the raw assembly template https://insillyclo.pasteur.cloud/assembly/8/
    - If more than 8 input plasmids, additional columns can be added directly in the template
    - Edit the restriction enzyme if other than BsaI
2. Prepare a *SequenceDB.zi*p archive with plasmids in Genbank format (only .gb files)
3. Fill in the template with genetic designs using the Genbank file names of the input plasmids
    - Since specification is made using file names, *iP_mapping* file can be omitted
4. Upload on Assembly Simulator module the filled template and the zip archive
5. Run the simulation and generate the campaign data

### Text S2: Standard typed cloning workflow

#### Initialization

1. Define the assembly on Assembly Designer or directly use a public assembly (https://insillyclo.pasteur.cloud/assembly/), and download the associated template file

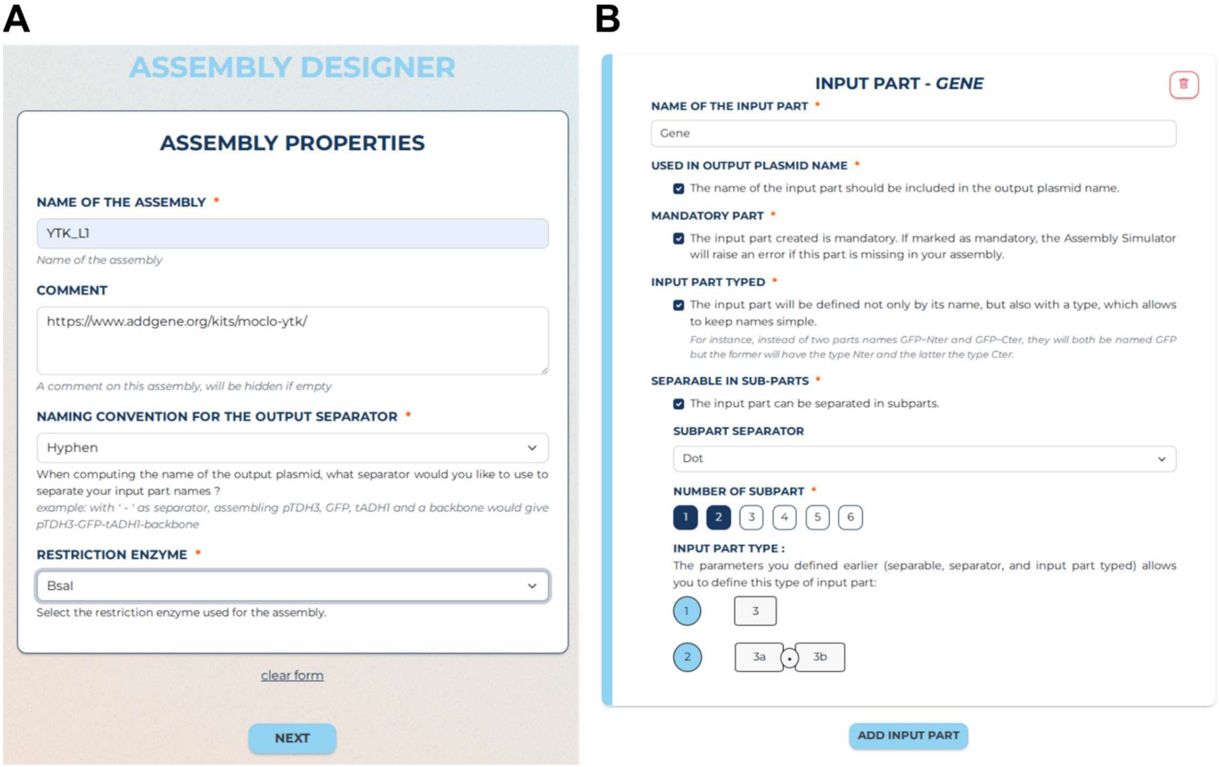


***Figure 1: Interface of the Assembly Designer module****. (A) Definition of the assembly properties. (B) Definition of the input part properties with the "Gene" example. Users can add as many input part as needed, and for each define their characteristics.*

- Types of parts and subparts cannot be edited in the web application. They are by default 1, 2 … for part and 1a, 1b… for subparts. Users need to modify them in the *template.xlsx* file once downloaded. Only numbers and letters are accepted.

2. Prepare lab data
   - *IP_mapping.csv* which maps genetic part names with Genbank file names (can be omitted if Genbank file names are used when specifying genetic designs).
   - *SequenceDB.zip* archive with the Genbank files (only .gb files).

**Campaign**

1. Fill in *template.xlsx* with genetic designs and save it as a campaign file
   - The type of each output plasmid is set manually in the "OutputType" column. For most MoClo system, it depends on the backbone used. Alternatively, it can depend on the connectors as in the YTK[1].
2. Upload on Assembly Simulator the filled template, then the lab data
3. Run the simulation to generate campaign data
   - Plasmid maps and input file of Auto-GG are generated by default
   - To compute dilutions, input plasmid concentrations are needed. They can be added in *IP_mapping.csv* from the beginning, or added in *input-plasmid-concentrations.csv* downloaded from the "Compute dilution" section of the result page
   - Upload the primer database if colony PCR is simulated

Additional information and example of datasets are available on https://insillyclo.pasteur.cloud/tutorial.

### Text S3: InSillyClo with other MoClo systems

Public assemblies for several MoClo kits are available on the web application (https://insillyclo.pasteur.cloud/assembly/). Additionally, we generated the input files (*template.xlsx, iP_maping.csv* and *SequenceDB.zip*) and a demo of campaign for some of them. When downloading *template.xlsx* from the web application, part types are numbers (1, 2 …) so users must change them if needed. We present below possible typing systems for each of these MoClo kits. They are all hierarchical so input and output types of each level are made compatible. The associated files are available at https://gitlab.pasteur.fr/insillyclo/insillyclo-web/-/tree/main/src/InSillyCloWeb/test_data/moclo_assembly_template?ref_type=heads.

#### MoClo Plant Part Kit[2]

This system allows typical Transcriptional Unit (TU) assembly in the level 1, followed by multi-Transcriptional Unit (mTU) assembly in level 2. Modularity of the TU structure is well captured by using subparts. For instance, the promoter region can be made one part (Pro5U), two subparts (Pro and 5U), or three subparts (Pro, 5Uf, NT1). This system allows the assembly of multiTU together through an iterative process with M and P levels. We think that "untyped" (ie raw) assemblies are more suited for these levels since designs become highly custom with the possibility to have any TU at the start or at the end of an assembly. In addition, standardized names of these multiTU plasmids can be rather long, making more convenient to use directly Genbank file names in design specification, as shown in Text S1. Of note, the first M assembly is similar to a Level 2 assembly.

| Assembly level | Input part types | Output part types |
|---|---|---|
| Level 1 | -Pro5U, [Pro, 5U], [Pro, 5Uf, NT1]<br>-CDS1, [SP, CDS2]<br>-CT<br>-3UTer, [3U, Ter]<br>-BL1 (Backbone L1) | TU1, TU2, TU3, TU4, TU5, TU6, TU7 |
| Level 2/M1 | -TU1<br>-TU2<br>-TU3<br>-TU4<br>-TU5<br>-TU6<br>-TU7<br>-ELL2/ELLM (End-Linker L2/M)<br>-BL2/BLM (Backbone L2/M) | mTU |

*Table 1: Typing system for the Plant Part Kit Moclo.*

**EcoFlex Moclo**[3]

EcoFlex has the typical Level 1 and Level 2 assemblies for construction of TU and mTU. As the Plant Part Toolkit, it allows to assemble together several mTUs, with the difference of being a unique additional assembly (i.e. Level 3 instead of Level M/P) with a simpler typing system.

| Assembly level | Input part types | Output part types |
|---|---|---|
| Level 1 | -1, [1a, 1b], [1a, 1b1, 1b2] (promoter, RBS and signal peptide)<br>-2 (CDS)<br>-3 (Terminator)<br>-BL1 (Backbone L1) | TUA, TUB, TUC, TUD, TUE |
| Level 2 | -TUA<br>-TUB<br>-TUC<br>-TUD<br>-TUE<br>-BL2 (Backbone L2) | mTUA, mTUB, mTUC, mTUD |
| Level 3 | -mTUA<br>-mTUB<br>-mTUC<br>-mTUD<br>-BL3 (Backbone L3) | mTU |

*Table 2: Typing system for the EcoFlex Moclo.*

**Cidar Moclo**[4]

This Moclo kit is similar than the EcoFlex in that it allows 3 level of assemblies but the level 0 design is different with promoters and terminator containing TU-specific overhang. Another difference is the reuse of the level 1 backbone in the level 3.

| Assembly level | Input part types | Output part types |
|---|---|---|
| Level 1 | -1 (promoter)<br>-2 (RBS)<br>-3 (CDS)<br>-4 (Terminator)<br>-BK (Backbone Kanamycin) | TUKae, TUKef, TUKfg, TUKgh |
| Level 2 | -TUKae<br>-TUKef<br>-TUKfg<br>-TUKgh<br>-BA (Backbone Ampicillin) | TUAae, TUAef, TUAfg, TUAgh |

| | | |
|---|---|---|
| Level 3 | -TUAae<br>-TUAef<br>-TUAfg<br>-TUAgh<br>-BK (Backbone Kanamycin) | mTU |

*Table 3: Typing system for the Cidar Moclo.*

### Text S4: Local implementation with the command line tool

We describe below the workflow taking as example the yeast display campaign. The full documentation is available at https://insillyclo.pages.pasteur.fr/insillyclo-cli/ and test datasets can be found at https://gitlab.pasteur.fr/insillyclo/insillyclo-web/-/tree/main/src/InSillyCloWeb/test_data/tutorial/Cli_assembly?ref_type=heads.

#### Setup

1. Get a local implementation of python
   - Download miniforge at https://conda-forge.org/miniforge/, and keep the "create shortcut" option during the installation.
2. Open miniforge prompt (from the start menu in Windows)
3. Run `pip install insillyclo`

#### *Template* command for the initialization

1. Run the *template* command

```
insillyclo template C:\Users\Henri\Documents\InSillyClo\Template_YTK_L1_BASE.xlsx
--name "YTK_L1"
--enzyme "BsaI"
--separator -
--input-part ConL
--input-part Promoter
--input-part Gene
--input-part Terminator
--input-part ConR
--input-part Backbone
```

Note: it needs to be given as a single line

2. If needed, edit the template file in a spreadsheet program to adapt input part parameters

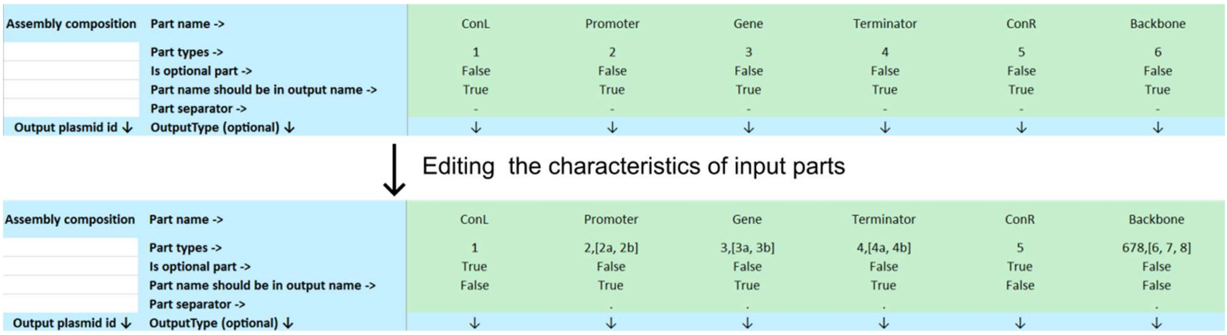


*Figure 2: Template editing with a spreadsheet program.*

- Part types are modified to add sub-parts and to correspond the grammar of the Moclo system.
- ConL and ConR are made optional.
- ConL, ConR and Backbone are removed from the output name.
- Part separator is set to be the dot "." or none.

3. Prepare lab data (*iP_mapping.csv*, plasmid repository, and table of primers)

As opposed to *SequenceDB.zip* in the web application, the plasmid repository is the folder containing the Genbank files (only .gb). It does not need to be zipped.

***Simulate*** **command for the campaign**

1. Fill in the template with genetic designs
2. Run the *simulate* command

```
insillyclo simulate
--input-template-filled C:\Users\Henri\Documents\InSillyClo\Campaign_YTK_display_cli.xlsx
--input-parts-file C:\Users\Henri\Documents\InSillyClo\iP_mapping_YTK.csv
--plasmid-repository C:\Users\Henri\Documents\InSillyClo\SequenceDB_YTK
--output-dir C:\Users\Henri\Documents\InSillyClo\CampaignData_display_cli
--restriction-enzyme-gel NotI
--primer-pair P29,P30
--primers-file C:\Users\Henri\Documents\InSillyClo\DB_primer.csv
--default-mass-concentration 200
```

Users of the command line tool need to specify all the results they want directly in the *simulate* command. It is possible to simulate PCR with several primer pairs, and digestion with several restriction enzymes (see details at https://insillyclo.pages.pasteur.fr/insillyclo-cli/simulate.html#insillyclo-simulate).

For users reluctant to upload their files on the server in the Assembly Simulator module, it is still possible to combine the web application and the command line tool. Concretely, the Assembly Designer module is used to generate the *template.xlsx* file, followed by the *simulate* command of the command line tool which handles files locally.

### File S1: Campaign_display_Raw.xlsx

| Assembly settings | |
|---|---|
| Restriction enzyme | BsaI |
| Name | Raw_BsaI |
| Output separator | - |

| Assembly composition | Part name -> | iP1 | iP2 | iP3 | iP4 | iP5 | iP6 | iP7 | iP8 |
|---|---|---|---|---|---|---|---|---|---|
| | Part types -> | | | | | | | | |
| | Is optional part -> | True | True | True | True | True | True | True | True |
| | Part name should be in output name -> | True | True | True | True | True | True | True | True |
| | Part separator -> | | | | | | | | |
| Output plasmid id ↓ | OutputType (optional) ↓ | ↓ | ↓ | ↓ | ↓ | ↓ | ↓ | ↓ | ↓ |
| pIB001 | | ConLS | pTET | SPsuc2 | EGII | AGA2 | tTDH1_4b | ConR1 | AmpR |
| pIB002 | | ConLS | pTET | SPplb1 | EGII | AGA2 | tTDH1_4b | ConR1 | AmpR |
| pIB003 | | ConLS | pTET | SPksh1 | EGII | AGA2 | tTDH1_4b | ConR1 | AmpR |
| pIB004 | | ConLS | pTET | SPsrl1 | EGII | AGA2 | tTDH1_4b | ConR1 | AmpR |
| pIB005 | | ConLS | pTET | SPcrh1 | EGII | AGA2 | tTDH1_4b | ConR1 | AmpR |
| pIB006 | | ConLS | pTET | SPost1p | EGII | AGA2 | tTDH1_4b | ConR1 | AmpR |
| pIB007 | | ConLS | pTET | SPsuc2 | EGII | SED1 | tTDH1_4b | ConR1 | AmpR |
| pIB008 | | ConLS | pTET | SPplb1 | EGII | SED1 | tTDH1_4b | ConR1 | AmpR |
| pIB009 | | ConLS | pTET | SPksh1 | EGII | SED1 | tTDH1_4b | ConR1 | AmpR |
| pIB010 | | ConLS | pTET | SPsrl1 | EGII | SED1 | tTDH1_4b | ConR1 | AmpR |
| pIB011 | | ConLS | pTET | SPcrh1 | EGII | SED1 | tTDH1_4b | ConR1 | AmpR |
| pIB012 | | ConLS | pTET | SPost1p | EGII | SED1 | tTDH1_4b | ConR1 | AmpR |
| pIB013 | | ConLS | pTET | SPsuc2 | EGII | TIP1 | tTDH1_4b | ConR1 | AmpR |
| pIB014 | | ConLS | pTET | SPplb1 | EGII | TIP1 | tTDH1_4b | ConR1 | AmpR |
| pIB015 | | ConLS | pTET | SPksh1 | EGII | TIP1 | tTDH1_4b | ConR1 | AmpR |
| pIB016 | | ConLS | pTET | SPsrl1 | EGII | TIP1 | tTDH1_4b | ConR1 | AmpR |
| pIB017 | | ConLS | pTET | SPcrh1 | EGII | TIP1 | tTDH1_4b | ConR1 | AmpR |
| pIB018 | | ConLS | pTET | SPost1p | EGII | TIP1 | tTDH1_4b | ConR1 | AmpR |
| pIB019 | | ConLS | pTET | SPsuc2 | EGII | CWP2 | tTDH1_4b | ConR1 | AmpR |
| pIB020 | | ConLS | pTET | SPplb1 | EGII | CWP2 | tTDH1_4b | ConR1 | AmpR |
| pIB021 | | ConLS | pTET | SPksh1 | EGII | CWP2 | tTDH1_4b | ConR1 | AmpR |
| pIB022 | | ConLS | pTET | SPsrl1 | EGII | CWP2 | tTDH1_4b | ConR1 | AmpR |
| pIB023 | | ConLS | pTET | SPcrh1 | EGII | CWP2 | tTDH1_4b | ConR1 | AmpR |
| pIB024 | | ConLS | pTET | SPost1p | EGII | CWP2 | tTDH1_4b | ConR1 | AmpR |
| pIB025 | | ConL1 | pTET | SPsuc2 | CBHI | AGA2 | tTDH1_4b | ConR2 | AmpR |
| pIB026 | | ConL1 | pTET | SPplb1 | CBHI | AGA2 | tTDH1_4b | ConR2 | AmpR |
| pIB027 | | ConL1 | pTET | SPksh1 | CBHI | AGA2 | tTDH1_4b | ConR2 | AmpR |
| pIB028 | | ConL1 | pTET | SPsrl1 | CBHI | AGA2 | tTDH1_4b | ConR2 | AmpR |
| pIB029 | | ConL1 | pTET | SPcrh1 | CBHI | AGA2 | tTDH1_4b | ConR2 | AmpR |
| pIB030 | | ConL1 | pTET | SPost1p | CBHI | AGA2 | tTDH1_4b | ConR2 | AmpR |
| pIB031 | | ConL1 | pTET | SPsuc2 | CBHI | SED1 | tTDH1_4b | ConR2 | AmpR |
| pIB032 | | ConL1 | pTET | SPplb1 | CBHI | SED1 | tTDH1_4b | ConR2 | AmpR |
| pIB033 | | ConL1 | pTET | SPksh1 | CBHI | SED1 | tTDH1_4b | ConR2 | AmpR |
| pIB034 | | ConL1 | pTET | SPsrl1 | CBHI | SED1 | tTDH1_4b | ConR2 | AmpR |
| pIB035 | | ConL1 | pTET | SPcrh1 | CBHI | SED1 | tTDH1_4b | ConR2 | AmpR |
| pIB036 | | ConL1 | pTET | SPost1p | CBHI | SED1 | tTDH1_4b | ConR2 | AmpR |
| pIB037 | | ConL1 | pTET | SPsuc2 | CBHI | TIP1 | tTDH1_4b | ConR2 | AmpR |
| pIB038 | | ConL1 | pTET | SPplb1 | CBHI | TIP1 | tTDH1_4b | ConR2 | AmpR |
| pIB039 | | ConL1 | pTET | SPksh1 | CBHI | TIP1 | tTDH1_4b | ConR2 | AmpR |
| pIB040 | | ConL1 | pTET | SPsrl1 | CBHI | TIP1 | tTDH1_4b | ConR2 | AmpR |
| pIB041 | | ConL1 | pTET | SPcrh1 | CBHI | TIP1 | tTDH1_4b | ConR2 | AmpR |
| pIB042 | | ConL1 | pTET | SPost1p | CBHI | TIP1 | tTDH1_4b | ConR2 | AmpR |
| pIB043 | | ConL1 | pTET | SPsuc2 | CBHI | CWP2 | tTDH1_4b | ConR2 | AmpR |

| | | | | | | | | | |
|---|---|---|---|---|---|---|---|---|---|
| pIB044 | | ConL1 | pTET | SPplb1 | CBHI | CWP2 | tTDH1_4b | ConR2 | AmpR |
| pIB045 | | ConL1 | pTET | SPksh1 | CBHI | CWP2 | tTDH1_4b | ConR2 | AmpR |
| pIB046 | | ConL1 | pTET | SPsrl1 | CBHI | CWP2 | tTDH1_4b | ConR2 | AmpR |
| pIB047 | | ConL1 | pTET | SPcrh1 | CBHI | CWP2 | tTDH1_4b | ConR2 | AmpR |
| pIB048 | | ConL1 | pTET | SPost1p | CBHI | CWP2 | tTDH1_4b | ConR2 | AmpR |
| pIB049 | | ConL2 | pTET | SPsuc2 | BGL | AGA2 | tTDH1_4b | ConR3 | AmpR |
| pIB050 | | ConL2 | pTET | SPplb1 | BGL | AGA2 | tTDH1_4b | ConR3 | AmpR |
| pIB051 | | ConL2 | pTET | SPksh1 | BGL | AGA2 | tTDH1_4b | ConR3 | AmpR |
| pIB052 | | ConL2 | pTET | SPsrl1 | BGL | AGA2 | tTDH1_4b | ConR3 | AmpR |
| pIB053 | | ConL2 | pTET | SPcrh1 | BGL | AGA2 | tTDH1_4b | ConR3 | AmpR |
| pIB054 | | ConL2 | pTET | SPost1p | BGL | AGA2 | tTDH1_4b | ConR3 | AmpR |
| pIB055 | | ConL2 | pTET | SPsuc2 | BGL | SED1 | tTDH1_4b | ConR3 | AmpR |
| pIB056 | | ConL2 | pTET | SPplb1 | BGL | SED1 | tTDH1_4b | ConR3 | AmpR |
| pIB057 | | ConL2 | pTET | SPksh1 | BGL | SED1 | tTDH1_4b | ConR3 | AmpR |
| pIB058 | | ConL2 | pTET | SPsrl1 | BGL | SED1 | tTDH1_4b | ConR3 | AmpR |
| pIB059 | | ConL2 | pTET | SPcrh1 | BGL | SED1 | tTDH1_4b | ConR3 | AmpR |
| pIB060 | | ConL2 | pTET | SPost1p | BGL | SED1 | tTDH1_4b | ConR3 | AmpR |
| pIB061 | | ConL2 | pTET | SPsuc2 | BGL | TIP1 | tTDH1_4b | ConR3 | AmpR |
| pIB062 | | ConL2 | pTET | SPplb1 | BGL | TIP1 | tTDH1_4b | ConR3 | AmpR |
| pIB063 | | ConL2 | pTET | SPksh1 | BGL | TIP1 | tTDH1_4b | ConR3 | AmpR |
| pIB064 | | ConL2 | pTET | SPsrl1 | BGL | TIP1 | tTDH1_4b | ConR3 | AmpR |
| pIB065 | | ConL2 | pTET | SPcrh1 | BGL | TIP1 | tTDH1_4b | ConR3 | AmpR |
| pIB066 | | ConL2 | pTET | SPost1p | BGL | TIP1 | tTDH1_4b | ConR3 | AmpR |
| pIB067 | | ConL2 | pTET | SPsuc2 | BGL | CWP2 | tTDH1_4b | ConR3 | AmpR |
| pIB068 | | ConL2 | pTET | SPplb1 | BGL | CWP2 | tTDH1_4b | ConR3 | AmpR |
| pIB069 | | ConL2 | pTET | SPksh1 | BGL | CWP2 | tTDH1_4b | ConR3 | AmpR |
| pIB070 | | ConL2 | pTET | SPsrl1 | BGL | CWP2 | tTDH1_4b | ConR3 | AmpR |
| pIB071 | | ConL2 | pTET | SPcrh1 | BGL | CWP2 | tTDH1_4b | ConR3 | AmpR |
| pIB072 | | ConL2 | pTET | SPost1p | BGL | CWP2 | tTDH1_4b | ConR3 | AmpR |
| pIB073 | | ConL3 | pCCW12 | AGA1 | tPGK1_4 | ConR4 | AmpR | | |
| pIB074 | | ConL4 | pRAD27 | rtTA | tENO2_4 | ConR5 | AmpR | | |
| pIB075 | | ConL5 | pUPR | Venus_3 | tADH1_4 | ConRE | AmpR | | |

### File S2: IP_mapping_Raw.csv

| pID | Name |
|---|---|
| pYTK002 | ConLS |
| pYTK003 | ConL1 |
| pYTK004 | ConL2 |
| pYTK005 | ConL3 |
| pYTK006 | ConL4 |
| pYTK007 | ConL5 |
| pYTK008 | ConLS' |
| pYTK009 | pTDH3 |
| pYTK010 | pCCW12 |
| pYTK011 | pPGK1 |
| pYTK012 | pHHF2 |
| pYTK013 | pTEF1 |
| pYTK014 | pTEF2 |
| pYTK015 | pHHF1 |
| pYTK016 | pHTB2 |
| pYTK017 | pRPL18B |
| pYTK018 | pALD6 |
| pYTK019 | pPAB1 |
| pYTK020 | pRET2 |
| pYTK021 | pRNR1 |
| pYTK022 | pSAC6 |
| pYTK023 | pRNR2 |
| pYTK024 | pPOP6 |
| pYTK025 | pRAD27 |
| pYTK026 | pPSP2 |
| pYTK027 | pREV1 |
| pYTK028 | pMFA1 |
| pYTK029 | pMFalpha2 |
| pYTK030 | pGAL1 |
| pYTK031 | pCUP1 |
| pYTK032 | mTurquoise2_3 |
| pYTK033 | Venus_3 |
| pYTK034 | mRuby2_3 |
| pYTK035 | I-SceI |
| pYTK036 | Cas9 |
| pYTK037 | mTurquoise2_3a |
| pYTK038 | Venus_3a |
| pYTK039 | mRuby2_3a |
| pYTK040 | 3xFLAG-6xHIS_3a |
| pYTK041 | UbiM |
| pYTK042 | UbiY |
| pYTK043 | UbiR |
| pYTK044 | mTurquoise2_3b |
| pYTK045 | Venus_3b |
| pYTK046 | mRuby2_3b |
| pYTK047 | GFP_dropout |
| pYTK048 | Spacer |
| pYTK049 | I-SceI reco site |
| pYTK050 | sgRNA dropout |
| pYTK051 | tENO1_4 |
| pYTK052 | tSSA1_4 |
| pYTK053 | tADH1_4 |
| pYTK054 | tPGK1_4 |
| pYTK055 | tENO2_4 |
| pYTK056 | tTDH1_4 |
| pYTK057 | mTurquoise2_4a |
| pYTK058 | Venus_4a |
| pYTK059 | mRuby2_4a |
| pYTK060 | 3xFLAG~6xHIS_4a |

| pID | Name |
|---|---|
| pMYT002 | pLacD |
| pMYT003 | pLacE |
| pMYT004 | pLacC |
| pMYT005 | pLacFEC |
| pMYT006 | pLacHTA1 |
| pMYT007 | pLacFBA1 |
| pMYT008 | pLacENO2 |
| pMYT009 | pLacSpTDH3 |
| pMYT010 | pTET2 |
| pMYT011 | pTET3 |
| pMYT012 | pTET4 |
| pMYT013 | pTET5 |
| pMYT014 | pTET6 |
| pMYT015 | pTET |
| pMYT016 | Z3EV |
| pMYT017 | LacI |
| pMYT018 | rtTA |
| pMYT019 | mScarletI |
| pMYT020 | mNeonGreen |
| pMYT021 | mTagBFP2 |
| pMYT022 | tTDH3 |
| pMYT023 | tTEF1 |
| pMYT024 | tTDH2 |
| pMYT025 | tPDC1 |
| pMYT039 | AmpRS1 |
| pMYT040 | AmpR1E |
| pMYT041 | AmpR12 |
| pMYT042 | AmpR2E |
| pMYT043 | AmpR23 |
| pMYT044 | AmpR3E |
| pMYT045 | AmpR34 |
| pMYT046 | AmpR4E |
| pMYT047 | AmpR45 |
| pMYT048 | AmpR5E |
| pMYT049 | AmpR56 |
| pMYT050 | AmpR6E |
| pMYT051 | AmpR67 |
| pMYT052 | AmpR7E |
| pMYT053 | AmpR78 |
| pMYT054 | AmpR8E |
| pMYT055 | AmpR89 |
| pMYT056 | AmpR9E |
| pMYT057 | Spacer_TU1 |
| pMYT058 | SpacerE_TU2 |
| pMYT059 | Spacer_TU2 |
| pMYT060 | SpacerE_TU3 |
| pMYT061 | Spacer_TU3 |
| pMYT062 | SpacerE_TU4 |
| pMYT063 | Spacer_TU4 |
| pMYT064 | SpacerE_TU5 |
| pMYT065 | Spacer_TU5 |
| pMYT066 | SpacerE_TU6 |
| pMYT067 | Spacer_TU6 |
| pMYT068 | SpacerE_TU7 |
| pMYT069 | Spacer_TU7 |
| pMYT070 | SpacerE_TU8 |
| pMYT071 | Spacer_TU8 |
| pMYT072 | SpacerE_TU9 |
| pMYT073 | Spacer_TU9 |

| | |
|---|---|
| pYTK061 | tENO1_4b |
| pYTK062 | tSSA1_4b |
| pYTK063 | tADH1_4b |
| pYTK064 | tPGK1_4b |
| pYTK065 | tENO2_4b |
| pYTK066 | tTDH1_4b |
| pYTK067 | ConR1 |
| pYTK068 | ConR2 |
| pYTK069 | ConR3 |
| pYTK070 | ConR4 |
| pYTK071 | ConR5 |
| pYTK072 | ConRE |
| pYTK073 | ConRE' |
| pYTK074 | URA3marker |
| pYTK075 | LEU2marker |
| pYTK076 | HIS3marker |
| pYTK077 | KanamycinR |
| pYTK078 | NourseothricinR |
| pYTK079 | HygromycinR |
| pYTK080 | ZeocinR |
| pYTK081 | CEN6/ARS4 |
| pYTK082 | 2micron |
| pYTK083 | AmpR_8 |
| pYTK084 | KanR_8 |
| pYTK085 | SpecR_8 |
| pYTK086 | URA3 3' |
| pYTK087 | LEU2 3' |
| pYTK088 | HO 3' |
| pYTK089 | AmpR_8a |
| pYTK090 | KanR_8a |
| pYTK091 | SpecR_8a |
| pYTK092 | URA3 5' |
| pYTK093 | LEU2 5' |
| pYTK094 | HO 5' |
| pYTK095 | AmpR |
| pYTK096 | Integ URA3 |
| pUPR066 | pUPR |
| pYSD001 | SPaga2 |
| pYSD002 | SPcrh1 |
| pYSD003 | SPecm14 |
| pYSD004 | SPsuc2 |
| pYSD005 | SPksh1 |
| pYSD006 | SPmfalpha1p |
| pYSD007 | SPost1p |
| pYSD008 | SPplb1 |
| pYSD009 | SPsrl1 |
| pYSD021 | SPmfalpha1pp8 |
| pYSD022 | SPmfalpha1opt |
| pYSD023 | SPost1mfalpha1 |
| pYSD024 | SPmfalpha1WT |
| pYSD041 | YAR066W |
| pYSD042 | HSP150 |
| pYSD043 | SCW4(111) |
| pYSD044 | SCW4(142) |
| pYSD061 | SUC2 |
| pYSD063 | mRuby2 |
| pYSD080 | AGA1 |
| pYSD081 | AGA2 |
| pYSD082 | CWP2 |
| pYSD083 | SED1 |
| pYSD084 | TIP1 |
| pYSD085 | 649stalk |
| pMYT001 | pZ3 |

| | |
|---|---|
| pMYT074 | SpacerE_TU10 |
| pMYT075 | BBInt1 |
| pMYT076 | BBInt2 |
| pMYT077 | BBInt3 |
| pMYT078 | BBInt4 |
| pMYT079 | BBInt5 |
| pMYT080 | BBInt6 |
| pMYT081 | BBInt7 |
| pMYT082 | BBInt8 |
| pMYT083 | BBInt9 |
| pMYT084 | BBInt10 |
| pMYT085 | SpacerInt1 |
| pMYT086 | SpacerInt2 |
| pMYT087 | SpacerInt3 |
| pMYT088 | SpacerInt4 |
| pMYT089 | SpacerInt5 |
| pMYT090 | SpacerInt6 |
| pMYT091 | SpacerInt7 |
| pMYT092 | SpacerInt8 |
| pMYT093 | SpacerInt9 |
| pMYT094 | SpacerInt10 |
| pCDE067 | EGII |
| pCDE068 | BGL |
| pCDE069 | CBHI |

### File S3: Campaign_display_L1.xlsx

| Assembly settings | |
|---|---|
| Restriction enzyme | BsaI |
| Name | YTK_L1 |
| Output separator | - |

| Assembly composition | Part name -> | ConL | Promoter | Gene | Terminator | ConR | Backbone |
|---|---|---|---|---|---|---|---|
| | Part types -> | 1 | 2 | 3,[3a, 3b] | 4,[4a, 4b] | 5 | 678,[6, 7, 8] |
| | Is optional part -> | True | False | False | False | True | False |
| | Part name should be in output name -> | False | True | True | True | False | False |
| | Part separator -> | | . | . | . | | . |
| Output plasmid id ↓ | OutputType (optional) ↓ | ↓ | ↓ | ↓ | ↓ | ↓ | ↓ |
| pIB001 | TU1 | ConLS | pTET | SPsuc2.EGII | AGA2.tTDH1 | ConR1 | AmpR |
| pIB002 | TU1 | ConLS | pTET | SPplb1.EGII | AGA2.tTDH1 | ConR1 | AmpR |
| pIB003 | TU1 | ConLS | pTET | SPksh1.EGII | AGA2.tTDH1 | ConR1 | AmpR |
| pIB004 | TU1 | ConLS | pTET | SPsrl1.EGII | AGA2.tTDH1 | ConR1 | AmpR |
| pIB005 | TU1 | ConLS | pTET | SPcrh1.EGII | AGA2.tTDH1 | ConR1 | AmpR |
| pIB006 | TU1 | ConLS | pTET | SPost1p.EGII | AGA2.tTDH1 | ConR1 | AmpR |
| pIB007 | TU1 | ConLS | pTET | SPsuc2.EGII | SED1.tTDH1 | ConR1 | AmpR |
| pIB008 | TU1 | ConLS | pTET | SPplb1.EGII | SED1.tTDH1 | ConR1 | AmpR |
| pIB009 | TU1 | ConLS | pTET | SPksh1.EGII | SED1.tTDH1 | ConR1 | AmpR |
| pIB010 | TU1 | ConLS | pTET | SPsrl1.EGII | SED1.tTDH1 | ConR1 | AmpR |
| pIB011 | TU1 | ConLS | pTET | SPcrh1.EGII | SED1.tTDH1 | ConR1 | AmpR |
| pIB012 | TU1 | ConLS | pTET | SPost1p.EGII | SED1.tTDH1 | ConR1 | AmpR |
| pIB013 | TU1 | ConLS | pTET | SPsuc2.EGII | TIP1.tTDH1 | ConR1 | AmpR |
| pIB014 | TU1 | ConLS | pTET | SPplb1.EGII | TIP1.tTDH1 | ConR1 | AmpR |
| pIB015 | TU1 | ConLS | pTET | SPksh1.EGII | TIP1.tTDH1 | ConR1 | AmpR |
| pIB016 | TU1 | ConLS | pTET | SPsrl1.EGII | TIP1.tTDH1 | ConR1 | AmpR |
| pIB017 | TU1 | ConLS | pTET | SPcrh1.EGII | TIP1.tTDH1 | ConR1 | AmpR |
| pIB018 | TU1 | ConLS | pTET | SPost1p.EGII | TIP1.tTDH1 | ConR1 | AmpR |
| pIB019 | TU1 | ConLS | pTET | SPsuc2.EGII | CWP2.tTDH1 | ConR1 | AmpR |
| pIB020 | TU1 | ConLS | pTET | SPplb1.EGII | CWP2.tTDH1 | ConR1 | AmpR |
| pIB021 | TU1 | ConLS | pTET | SPksh1.EGII | CWP2.tTDH1 | ConR1 | AmpR |
| pIB022 | TU1 | ConLS | pTET | SPsrl1.EGII | CWP2.tTDH1 | ConR1 | AmpR |
| pIB023 | TU1 | ConLS | pTET | SPcrh1.EGII | CWP2.tTDH1 | ConR1 | AmpR |
| pIB024 | TU1 | ConLS | pTET | SPost1p.EGII | CWP2.tTDH1 | ConR1 | AmpR |
| pIB025 | TU2 | ConL1 | pTET | SPsuc2.CBHI | AGA2.tTDH1 | ConR2 | AmpR |
| pIB026 | TU2 | ConL1 | pTET | SPplb1.CBHI | AGA2.tTDH1 | ConR2 | AmpR |
| pIB027 | TU2 | ConL1 | pTET | SPksh1.CBHI | AGA2.tTDH1 | ConR2 | AmpR |
| pIB028 | TU2 | ConL1 | pTET | SPsrl1.CBHI | AGA2.tTDH1 | ConR2 | AmpR |
| pIB029 | TU2 | ConL1 | pTET | SPcrh1.CBHI | AGA2.tTDH1 | ConR2 | AmpR |
| pIB030 | TU2 | ConL1 | pTET | SPost1p.CBHI | AGA2.tTDH1 | ConR2 | AmpR |
| pIB031 | TU2 | ConL1 | pTET | SPsuc2.CBHI | SED1.tTDH1 | ConR2 | AmpR |

| | | | | | | | |
|---|---|---|---|---|---|---|---|
| pIB032 | TU2 | ConL1 | pTET | SPplb1.CBHI | SED1.tTDH1 | ConR2 | AmpR |
| pIB033 | TU2 | ConL1 | pTET | SPksh1.CBHI | SED1.tTDH1 | ConR2 | AmpR |
| pIB034 | TU2 | ConL1 | pTET | SPsrl1.CBHI | SED1.tTDH1 | ConR2 | AmpR |
| pIB035 | TU2 | ConL1 | pTET | SPcrh1.CBHI | SED1.tTDH1 | ConR2 | AmpR |
| pIB036 | TU2 | ConL1 | pTET | SPost1p.CBHI | SED1.tTDH1 | ConR2 | AmpR |
| pIB037 | TU2 | ConL1 | pTET | SPsuc2.CBHI | TIP1.tTDH1 | ConR2 | AmpR |
| pIB038 | TU2 | ConL1 | pTET | SPplb1.CBHI | TIP1.tTDH1 | ConR2 | AmpR |
| pIB039 | TU2 | ConL1 | pTET | SPksh1.CBHI | TIP1.tTDH1 | ConR2 | AmpR |
| pIB040 | TU2 | ConL1 | pTET | SPsrl1.CBHI | TIP1.tTDH1 | ConR2 | AmpR |
| pIB041 | TU2 | ConL1 | pTET | SPcrh1.CBHI | TIP1.tTDH1 | ConR2 | AmpR |
| pIB042 | TU2 | ConL1 | pTET | SPost1p.CBHI | TIP1.tTDH1 | ConR2 | AmpR |
| pIB043 | TU2 | ConL1 | pTET | SPsuc2.CBHI | CWP2.tTDH1 | ConR2 | AmpR |
| pIB044 | TU2 | ConL1 | pTET | SPplb1.CBHI | CWP2.tTDH1 | ConR2 | AmpR |
| pIB045 | TU2 | ConL1 | pTET | SPksh1.CBHI | CWP2.tTDH1 | ConR2 | AmpR |
| pIB046 | TU2 | ConL1 | pTET | SPsrl1.CBHI | CWP2.tTDH1 | ConR2 | AmpR |
| pIB047 | TU2 | ConL1 | pTET | SPcrh1.CBHI | CWP2.tTDH1 | ConR2 | AmpR |
| pIB048 | TU2 | ConL1 | pTET | SPost1p.CBHI | CWP2.tTDH1 | ConR2 | AmpR |
| pIB049 | TU3 | ConL2 | pTET | SPsuc2.BGL | AGA2.tTDH1 | ConR3 | AmpR |
| pIB050 | TU3 | ConL2 | pTET | SPplb1.BGL | AGA2.tTDH1 | ConR3 | AmpR |
| pIB051 | TU3 | ConL2 | pTET | SPksh1.BGL | AGA2.tTDH1 | ConR3 | AmpR |
| pIB052 | TU3 | ConL2 | pTET | SPsrl1.BGL | AGA2.tTDH1 | ConR3 | AmpR |
| pIB053 | TU3 | ConL2 | pTET | SPcrh1.BGL | AGA2.tTDH1 | ConR3 | AmpR |
| pIB054 | TU3 | ConL2 | pTET | SPost1p.BGL | AGA2.tTDH1 | ConR3 | AmpR |
| pIB055 | TU3 | ConL2 | pTET | SPsuc2.BGL | SED1.tTDH1 | ConR3 | AmpR |
| pIB056 | TU3 | ConL2 | pTET | SPplb1.BGL | SED1.tTDH1 | ConR3 | AmpR |
| pIB057 | TU3 | ConL2 | pTET | SPksh1.BGL | SED1.tTDH1 | ConR3 | AmpR |
| pIB058 | TU3 | ConL2 | pTET | SPsrl1.BGL | SED1.tTDH1 | ConR3 | AmpR |
| pIB059 | TU3 | ConL2 | pTET | SPcrh1.BGL | SED1.tTDH1 | ConR3 | AmpR |
| pIB060 | TU3 | ConL2 | pTET | SPost1p.BGL | SED1.tTDH1 | ConR3 | AmpR |
| pIB061 | TU3 | ConL2 | pTET | SPsuc2.BGL | TIP1.tTDH1 | ConR3 | AmpR |
| pIB062 | TU3 | ConL2 | pTET | SPplb1.BGL | TIP1.tTDH1 | ConR3 | AmpR |
| pIB063 | TU3 | ConL2 | pTET | SPksh1.BGL | TIP1.tTDH1 | ConR3 | AmpR |
| pIB064 | TU3 | ConL2 | pTET | SPsrl1.BGL | TIP1.tTDH1 | ConR3 | AmpR |
| pIB065 | TU3 | ConL2 | pTET | SPcrh1.BGL | TIP1.tTDH1 | ConR3 | AmpR |
| pIB066 | TU3 | ConL2 | pTET | SPost1p.BGL | TIP1.tTDH1 | ConR3 | AmpR |
| pIB067 | TU3 | ConL2 | pTET | SPsuc2.BGL | CWP2.tTDH1 | ConR3 | AmpR |
| pIB068 | TU3 | ConL2 | pTET | SPplb1.BGL | CWP2.tTDH1 | ConR3 | AmpR |
| pIB069 | TU3 | ConL2 | pTET | SPksh1.BGL | CWP2.tTDH1 | ConR3 | AmpR |
| pIB070 | TU3 | ConL2 | pTET | SPsrl1.BGL | CWP2.tTDH1 | ConR3 | AmpR |
| pIB071 | TU3 | ConL2 | pTET | SPcrh1.BGL | CWP2.tTDH1 | ConR3 | AmpR |
| pIB072 | TU3 | ConL2 | pTET | SPost1p.BGL | CWP2.tTDH1 | ConR3 | AmpR |
| pIB073 | TU4 | ConL3 | pCCW12 | AGA1 | tPGK1 | ConR4 | AmpR |
| pIB074 | TU5 | ConL4 | pRAD27 | rtTA | tENO2 | ConR5 | AmpR |
| pIB075 | TU6 | ConL5 | pUPR | Venus | tADH1 | ConRE | AmpR |

### File S4: IP_mapping_YTK.csv

| pID | Name | Type |
|---|---|---|
| pYTK002 | ConLS | 1 |
| pYTK003 | ConL1 | 1 |
| pYTK004 | ConL2 | 1 |
| pYTK005 | ConL3 | 1 |
| pYTK006 | ConL4 | 1 |
| pYTK007 | ConL5 | 1 |
| pYTK008 | ConLS' | 1 |
| pYTK009 | pTDH3 | 2 |
| pYTK010 | pCCW12 | 2 |
| pYTK011 | pPGK1 | 2 |
| pYTK012 | pHHF2 | 2 |
| pYTK013 | pTEF1 | 2 |
| pYTK014 | pTEF2 | 2 |
| pYTK015 | pHHF1 | 2 |
| pYTK016 | pHTB2 | 2 |
| pYTK017 | pRPL18B | 2 |
| pYTK018 | pALD6 | 2 |
| pYTK019 | pPAB1 | 2 |
| pYTK020 | pRET2 | 2 |
| pYTK021 | pRNR1 | 2 |
| pYTK022 | pSAC6 | 2 |
| pYTK023 | pRNR2 | 2 |
| pYTK024 | pPOP6 | 2 |
| pYTK025 | pRAD27 | 2 |
| pYTK026 | pPSP2 | 2 |
| pYTK027 | pREV1 | 2 |
| pYTK028 | pMFA1 | 2 |
| pYTK029 | pMFalpha2 | 2 |
| pYTK030 | pGAL1 | 2 |
| pYTK031 | pCUP1 | 2 |
| pYTK032 | mTurquoise2 | 3 |
| pYTK033 | Venus | 3 |
| pYTK034 | mRuby2 | 3 |
| pYTK035 | I-SceI | 3 |
| pYTK036 | Cas9 | 3 |
| pYTK037 | mTurquoise2 | 3a |
| pYTK038 | Venus | 3a |
| pYTK039 | mRuby2 | 3a |
| pYTK040 | 3xFLAG-6xHIS | 3a |
| pYTK041 | UbiM | 3a |
| pYTK042 | UbiY | 3a |
| pYTK043 | UbiR | 3a |
| pYTK044 | mTurquoise2 | 3b |
| pYTK045 | Venus | 3b |
| pYTK046 | mRuby2 | 3b |
| pYTK047 | GFP_dropout | 234 |
| pYTK048 | Spacer | 234 |
| pYTK049 | I-SceI reco site | 234 |
| pYTK050 | sgRNA dropout | 234 |
| pYTK051 | tENO1 | 4 |
| pYTK052 | tSSA1 | 4 |
| pYTK053 | tADH1 | 4 |
| pYTK054 | tPGK1 | 4 |
| pYTK055 | tENO2 | 4 |
| pYTK056 | tTDH1 | 4 |
| pYTK057 | mTurquoise2 | 4a |
| pYTK058 | Venus | 4a |
| pYTK059 | mRuby2 | 4a |
| pYTK060 | 3xFLAG~6xHIS | 4a |

| pID | Name | Type |
|---|---|---|
| pMYT002 | pLacD | 2 |
| pMYT003 | pLacE | 2 |
| pMYT004 | pLacC | 2 |
| pMYT005 | pLacFEC | 2 |
| pMYT006 | pLacHTA1 | 2 |
| pMYT007 | pLacFBA1 | 2 |
| pMYT008 | pLacENO2 | 2 |
| pMYT009 | pLacSpTDH3 | 2 |
| pMYT010 | pTET2 | 2 |
| pMYT011 | pTET3 | 2 |
| pMYT012 | pTET4 | 2 |
| pMYT013 | pTET5 | 2 |
| pMYT014 | pTET6 | 2 |
| pMYT015 | pTET | 2 |
| pMYT016 | Z3EV | 3 |
| pMYT017 | LacI | 3 |
| pMYT018 | rtTA | 3 |
| pMYT019 | mScarletI | 3 |
| pMYT020 | mNeonGreen | 3 |
| pMYT021 | mTagBFP2 | 3 |
| pMYT022 | tTDH3 | 4 |
| pMYT023 | tTEF1 | 4 |
| pMYT024 | tTDH2 | 4 |
| pMYT025 | tPDC1 | 4 |
| pMYT039 | AmpRS1 | 678 |
| pMYT040 | AmpR1E | 678 |
| pMYT041 | AmpR12 | 678 |
| pMYT042 | AmpR2E | 678 |
| pMYT043 | AmpR23 | 678 |
| pMYT044 | AmpR3E | 678 |
| pMYT045 | AmpR34 | 678 |
| pMYT046 | AmpR4E | 678 |
| pMYT047 | AmpR45 | 678 |
| pMYT048 | AmpR5E | 678 |
| pMYT049 | AmpR56 | 678 |
| pMYT050 | AmpR6E | 678 |
| pMYT051 | AmpR67 | 678 |
| pMYT052 | AmpR7E | 678 |
| pMYT053 | AmpR78 | 678 |
| pMYT054 | AmpR8E | 678 |
| pMYT055 | AmpR89 | 678 |
| pMYT056 | AmpR9E | 678 |
| pMYT057 | Spacer | TU1 |
| pMYT058 | SpacerE | TU2 |
| pMYT059 | Spacer | TU2 |
| pMYT060 | SpacerE | TU3 |
| pMYT061 | Spacer | TU3 |
| pMYT062 | SpacerE | TU4 |
| pMYT063 | Spacer | TU4 |
| pMYT064 | SpacerE | TU5 |
| pMYT065 | Spacer | TU5 |
| pMYT066 | SpacerE | TU6 |
| pMYT067 | Spacer | TU6 |
| pMYT068 | SpacerE | TU7 |
| pMYT069 | Spacer | TU7 |
| pMYT070 | SpacerE | TU8 |
| pMYT071 | Spacer | TU8 |
| pMYT072 | SpacerE | TU9 |
| pMYT073 | Spacer | TU9 |

| | | |
|---|---|---|
| pYTK061 | tENO1 | 4b |
| pYTK062 | tSSA1 | 4b |
| pYTK063 | tADH1 | 4b |
| pYTK064 | tPGK1 | 4b |
| pYTK065 | tENO2 | 4b |
| pYTK066 | tTDH1 | 4b |
| pYTK067 | ConR1 | 5 |
| pYTK068 | ConR2 | 5 |
| pYTK069 | ConR3 | 5 |
| pYTK070 | ConR4 | 5 |
| pYTK071 | ConR5 | 5 |
| pYTK072 | ConRE | 5 |
| pYTK073 | ConRE' | 5 |
| pYTK074 | URA3marker | 6 |
| pYTK075 | LEU2marker | 6 |
| pYTK076 | HIS3marker | 6 |
| pYTK077 | KanamycinR | 6 |
| pYTK078 | NourseothricinR | 6 |
| pYTK079 | HygromycinR | 6 |
| pYTK080 | ZeocinR | 6 |
| pYTK081 | CEN6/ARS4 | 7 |
| pYTK082 | 2micron | 7 |
| pYTK083 | AmpR | 8 |
| pYTK084 | KanR | 8 |
| pYTK085 | SpecR | 8 |
| pYTK086 | URA3 3' | 7 |
| pYTK087 | LEU2 3' | 7 |
| pYTK088 | HO 3' | 7 |
| pYTK089 | AmpR | 8a |
| pYTK090 | KanR | 8a |
| pYTK091 | SpecR | 8a |
| pYTK092 | URA3 5' | 8b |
| pYTK093 | LEU2 5' | 8b |
| pYTK094 | HO 5' | 8b |
| pYTK095 | AmpR | 678 |
| pYTK096 | B_URA3_SE | B |
| pUPR066 | pUPR | 2 |
| pYSD001 | SPaga2 | 3a |
| pYSD002 | SPcrh1 | 3a |
| pYSD003 | SPecm14 | 3a |
| pYSD004 | SPsuc2 | 3a |
| pYSD005 | SPksh1 | 3a |
| pYSD006 | SPmfalpha1p | 3a |
| pYSD007 | SPost1p | 3a |
| pYSD008 | SPplb1 | 3a |
| pYSD009 | SPsrl1 | 3a |
| pYSD021 | SPmfalpha1pp8 | 3a |
| pYSD022 | SPmfalpha1opt | 3a |
| pYSD023 | SPost1mfalpha1 | 3a |
| pYSD024 | SPmfalpha1WT | 3a |
| pYSD041 | YAR066W | 3a |
| pYSD042 | HSP150 | 3a |
| pYSD043 | SCW4(111) | 3a |
| pYSD044 | SCW4(142) | 3a |
| pYSD061 | SUC2 | 3b |
| pYSD063 | mRuby2 | 3b |
| pYSD080 | AGA1 | 3 |
| pYSD081 | AGA2 | 4a |
| pYSD082 | CWP2 | 4a |
| pYSD083 | SED1 | 4a |
| pYSD084 | TIP1 | 4a |
| pYSD085 | 649stalk | 4a |
| pMYT001 | pZ3 | 2 |

| | | |
|---|---|---|
| pMYT074 | SpacerE | TU10 |
| pMYT075 | B_SE_Int1 | B |
| pMYT076 | B_SE_Int2 | B |
| pMYT077 | B_SE_Int3 | B |
| pMYT078 | B_SE_Int4 | B |
| pMYT079 | B_SE_Int5 | B |
| pMYT080 | B_SE_Int6 | B |
| pMYT081 | B_SE_Int7 | B |
| pMYT082 | B_SE_Int8 | B |
| pMYT083 | B_SE_Int9 | B |
| pMYT084 | B_SE_Int10 | B |
| pMYT085 | SpacerInt1 | mTUinteg |
| pMYT086 | SpacerInt2 | mTUinteg |
| pMYT087 | SpacerInt3 | mTUinteg |
| pMYT088 | SpacerInt4 | mTUinteg |
| pMYT089 | SpacerInt5 | mTUinteg |
| pMYT090 | SpacerInt6 | mTUinteg |
| pMYT091 | SpacerInt7 | mTUinteg |
| pMYT092 | SpacerInt8 | mTUinteg |
| pMYT093 | SpacerInt9 | mTUinteg |
| pMYT094 | SpacerInt10 | mTUinteg |
| pCDE067 | EGII | 3b |
| pCDE068 | BGL | 3b |
| pCDE069 | CBHI | 3b |

### References


(1) Lee, M. E.; DeLoache, W. C.; Cervantes, B.; Dueber, J. E. A Highly Characterized Yeast Toolkit for Modular, Multipart Assembly. *ACS Synth. Biol.* **2015**, *4* (9), 975–986. https://doi.org/10.1021/sb500366v.

(2) Engler, C.; Youles, M.; Gruetzner, R.; Ehnert, T.-M.; Werner, S.; Jones, J. D. G.; Patron, N. J.; Marillonnet, S. A Golden Gate Modular Cloning Toolbox for Plants. *ACS Synth. Biol.* **2014**, *3* (11), 839–843. https://doi.org/10.1021/sb4001504.

(3) Moore, S. J.; Lai, H.-E.; Kelwick, R. J. R.; Chee, S. M.; Bell, D. J.; Polizzi, K. M.; Freemont, P. S. EcoFlex: A Multifunctional MoClo Kit for E. Coli Synthetic Biology. *ACS Synth. Biol.* **2016**, *5* (10), 1059–1069. https://doi.org/10.1021/acssynbio.6b00031.

(4) Iverson, S. V.; Haddock, T. L.; Beal, J.; Densmore, D. M. CIDAR MoClo: Improved MoClo Assembly Standard and New E. Coli Part Library Enable Rapid Combinatorial Design for Synthetic and Traditional Biology. *ACS Synth. Biol.* **2016**, *5* (1), 99–103. https://doi.org/10.1021/acssynbio.5b00124.
